# Maternal deprivation impairs social recognition by diminishing the number of adult-born granule cells in the dentate gyrus

**DOI:** 10.64898/2026.11.08.765070

**Authors:** Renée C. Waters, Isha R. Gore, Casey J. Brown, Emma J. Diethorn, Aimee L. Skweres-Gilmartin, Elizabeth Gould

## Abstract

Childhood neglect increases the likelihood of developing neuropsychiatric disease in adulthood. Maternal separation and early weaning (MSEW), a rodent model of maternal deprivation, was used to investigate effects on social recognition, a commonly impaired function in neuropsychiatric disorders in humans. We found that in adulthood, MSEW male mice showed social recognition impairments, and that MSEW female mice showed greater preference fo social novelty. MSEW male, but not female, mice had diminished numbers of adult-born granule cells (abGCs) in the dentate gyrus, as well as a lower density of abGC axons in the CA2 region compared to controls. MSEW males also showed a reduced rate of sharp wave ripples (SWRs) in the CA2 region in response to novel mouse exposure, as well as a reduction in differential SWR rates in response to novel and familiar mouse exposure. Living in an enriched environment restored the number of abGCs in the dentate gyrus of MSEW males to control levels and improved social recognition. Furthermore, chemogenetic activation of 4-6 week old, but not older, abGCs in a double transgenic model improved social recognition and partially restored CA2 SWR patterns in MSEW mice. These findings suggest that MSEW impairs social recognition through diminished adult neurogenesis, which can be restored in adulthood by living in a proneurogenic environment or by increasing the activity of existing abGCs.

## Introduction

Childhood maltreatment, including emotional and physical neglect, increases the likelihood of several neuropsychiatric diseases in adulthood (Felitti et al., 1998; Lippard and Nemeroff, 2020). Among the conditions linked to childhood neglect, anxiety disorders and clinical depression (Kuo et al., 2011; Merola et al., 2026) have been associated with deficits in social function (Ladegaard et al., 2014; Saris et al., 2017; Jin et al., 2023). Furthermore, among otherwise healthy individuals, those that were subjected to emotional neglect during childhood have deficits in face recognition in adulthood (Iffland and Neuner, 2020) suggesting an impairment in social memort. Studies have shown that emotional neglect during childhood is associated with smaller hippocampal volumes in adulthood (Frodl et al., 2010; Teicher and Samson, 2016). Since the hippocampus has been linked to social cognition (Montagrin et al., 2018; Schafer and Schiller, 2018; Leblanc and Ramirez, 2020) and has an extended postnatal development (Krogsrud et al., 2014), it may be especially vulnerable to the adverse effects of childhood neglect. Identifying neural mechanisms linking early life neglect to social processing is an important step in establishing effective interventions to reverse deficits and improve function. Toward this end, numerous studies have used animal models of neglect or deprivation, including maternal separation as well as maternal separation with early weaning, to explore neural and behavioral outcomes, with a strong focus on nonsocial behavior (Murthy et al., 2019; Bachiller et al., 2020; Talani et al., 2023; Wu et al., 2025). A few studies have shown that maternal separation diminishes social discrimination in mice (Franklin et al., 2011) and rats (Lukas et al., 2011; Kentrop et al., 2018).

Studies have shown that adult-born granule cells (abGCs) in the dentate gyrus support both nonsocial and social memory in mice (Jessberger et al., 2009; Suárez-Pereira and Carrion, 2015; Cope et al., 2020; Laham et al., 2024). While abGCs appear to influence nonsocial memory through their axonal connections with the CA3 region of the hippocampus (Gu et al., 2012; McHugh et al., 2022), abGCs support social memory through their connections with area CA2 (Laham et al., 2024). Chemogenetic inhibition of abGC projections to CA2, a region of the hippocampus important for social memory processing, impairs social recognition (Laham et al., 2024). Furthermore, inhibition of abGCs impairs the differential generation of CA2 sharp wave ripples (SWRs) in response to novel versus familiar mice (Laham et al., 2024). Coupled with evidence that CA2 SWRs are necessary for social recognition (Oliva et al., 2020), this raises the possibility that early life neglect impairs social recognition by producing a lasting change to adult neurogenesis in the dentate gyrus and hence its influence on the CA2 region. Some findings support this possibility by showing that maternal separation diminishes adult neurogenesis in rats (Mirescu et al., 2004; Hulsdorf et al., 2011), although this has not been explored in depth in mice. Additionally, the possibility that reduced adult neurogenesis is causally linked to behavioral outcomes of maternal separation has not yet been investigated.

To address these gaps, we used the maternal separation with early weaning (MSEW) paradigm to explore persistent effects in mice and found that adult males, but not females, showed social recognition deficits, accompanied by impaired adult neurogenesis, reduced abGC afferents to the CA2 region, and disrupted CA2 SWRs in response to social stimuli. We found that increasing the number of abGCs by enriched environment living reversed the negative effects of MSEW on social recognition, and that chemogenetic activation of the reduced number of abGCs in MSEW mice restored social recognition and CA2 network activity. These findings suggest that although early life deprivation produces sex-specific deficits in social cognition that last into adulthood, restoration of function can be established in adult males by environmental or chemogenetic approaches.

## Methods

### Animals

All animal procedures were approved by the Princeton University Institutional Animal Care and Use Committee and were in accordance with the National Research Council Guide for the Care and Use of Laboratory Animals. C57BL/6 mice were obtained from Jackson laboratories and were used for most studies. Nestin-CreERT2 and R26-LSL-hM3Dq-DREADD transgenic mice were obtained from Jackson labs and bred in the Princeton Neuroscience Institute as single or double transgenic mice for behavioral and electrophysiology experiments. Initial studies used mixed-sex groups; however, some investigations were male-only as they were follow-up studies based on sex-specific MSEW-induced behavioral impairment. Transgenic mice were genotyped by Transnetyx from ear punches at P15 using real-time PCR. Control mice were weaned from their dams at P21 and housed in same sex groups of 4-5 per cage. Single and double transgenic animals were group housed with mixed genotypes. Animals were housed in Optimice cages on a reverse 12:12 h light-dark cycle.

### Maternal separation and early weaning

C57, single transgenic nestin-Cre, and double transgenic nestin-Cre:Gq pups were randomly assigned to either control-rearing or MSEW group as previously described (Murthy et al., 2019; Laham et al., 2022). From P3-P6, MSEW mice were separated from their dam for 4 hours per day. From P7-P16, separations increased to 8 hours per day. During separations, dams were removed from the home cage, and each cage containing the pups was placed on top of a thermal heating blanket maintained at 34°C. Pups remained with their littermates for the entire period of separation after which the dam was returned to the home cage. Maternally separated pups were then weaned four days earlier than controls, at P17. For transgenic experiments, all litters contained both genotypes.

### Environment Enrichment Housing

C57 control-reared and MSEW adult male mice were housed in an enriched environment or in standard control cages in a satellite housing facility in the Princeton Neuroscience Institute for 6 weeks, starting at P60. Enriched environment housing took place in large Sterilite containers (34”x18”x12”). Food and water were provided ad libitum. In addition to the standard nesting material and bedding, rodent pet-quality toys made of wood and plastic, shelter structures, and running wheels were provided. All objects were removed from the container once a week and cleaned. New objects were rotated into the environment once a week for the 6-week period to maintain novelty.

### Behavioral analysis

The direct social interaction test was used (Laham et al., 2021) to determine whether adult mice were able to discriminate between a novel and familiar mouse of the same sex and age. Because healthy adult mice prefer novelty, social recognition is measured as a decrease in investigation time by the test mouse from trial 1 with the “novel” mouse to trial 2 with the “familiar” mouse (Moy et al., 2004). Adult C57, single transgenic (nestin-Cre), and double transgenic (nestin-Cre:Gq) underwent the direct social interaction test. Following habituation to the room and the testing arena, the mice were tested in a 2-trial test. In trial 1 (novel), the test mouse was placed in an open field box and a novel sex- and age-matched C57 stimulus mouse was introduced. The mice were allowed to interact for 5 minutes. After a 1-hour delay in their respective home cages, trial 2 (familiar) began in which the test mouse was placed back in the open field with the same mouse for 5 minutes. Social exposures were videotaped and the time the test mouse spent investigating (sniffing, close following, allogrooming with a distance of ≤2 cm) the familiar or novel mouse was measured.

### Drug treatments

To induce recombination in double transgenic mice starting on P60, nestin-Cre and nestin-Cre:Gq mice were administered intraperitoneal (i.p.) injections of tamoxifen (75 mg/kg) dissolved in corn oil every other day for 9 days (5 total injections). 4 weeks after the last tamoxifen injection, nestin-Cre and nestin-Cre:Gq mice underwent social recognition testing (with novel and familiar stimulus mice) twice, once after a vehicle i.p. injection and once after a CNO (5 mg/kg) i.p. injection 30 minutes before trial 1. The order of drug administration (vehicle or CNO) was counterbalanced across groups. Electrode-implanted mice underwent direct social interaction testing after receiving systemic vehicle or CNO administration following the same experimental design. Mice were given a 48-hour minimum rest period between vehicle and CNO tests.

### Surgical manipulations

Mice were deeply anesthetized with isoflurane (2-3%) and placed in a stereotaxic apparatus (Kopf) under a temperature-controlled thermal blanket for all surgeries. The head was leveled using bregma, lambda, and medial-lateral reference points before virus injection or electrode implantation. MSEW and control-reared double transgenic male mice were implanted unilaterally with a custom-made 4-wire electrode array (Microprobes) into the right hemisphere targeting dorsal CA2 (AP: -1.82, ML: ±2.15, DV: -1.67). 4 bone screws were implanted into the skull with 1 screw positioned on the contralateral hemisphere serving as ground. The ground wire was wrapped around the ground screw and covered with metallic paint to ensure maximum contact. Electrode implants were kept in place using Metabond cement (Parkell) followed by dental cement (Bosworth Trim). Two weeks after surgery, electrode-implanted mice received systemic injections of vehicle or CNO 30 minutes prior to electrophysiological recordings.

For the retrovirus experiment, a replication-deficient retroviral vector derived from the Moloney leukemia virus, designed to express GFP under the CAG promoter, was employed to selectively transduce abGCs, following previously published methods (Briones et al., 2021; Diethorn and Gould, 2023). To deliver the retrovirus, adult male mice (P60) underwent identical surgical preparation as described above. A microsyringe was used to deliver CAG-GFP retrovirus bilaterally to the dorsal dentate gyrus (1.0 μl per injection at 0.15 μl/min; AP: −2.0, ML: ±1.3, DV (suprapyramidal blade): −2.0, DV (infrapyramidal blade): −2.3). Mice were perfused 6 weeks post-injection to examine GFP-labeled axon terminals in the CA2 and CA3 regions.

### Electrophysiology

Local field potentials (LFPs) were recorded from control-reared and MSEW nestin-Cre:Gq mice with CA2 electrodes, using a tethered head stage (HST/16D Gen2, Plexon). After 5 minutes of habituation, the behavioral testing paradigm consisted of a 3-minute baseline in the testing arena followed by a 5-minute social interaction period with either a novel or familiar mouse. Vehicle or CNO was i.p. administered 30 minutes before testing. LFPs were recorded continuously throughout baseline and stimulus trials. The data were transmitted to a receiver (OmniPlex Hydrid eChassis, Plexon) and recorded using PlexControl software (Plexon).

### Electrophysiological analyses

#### Power and phase-amplitude coupling

Using a custom Python script, theta (4-12 Hz), slow gamma (25-50 Hz), mid-gamma (50-100 Hz), and fast gamma (100-130) oscillations were extracted from the LFP signal obtained during the baseline, novel, and familiar conspecific during periods of immobility as determined using BORIS (Laham and Zahn, 2023).

All theta and gamma recordings underwent a Hilbert transform, and theta phase and gamma analytic signal were stored. Phase-amplitude coupling (PAC) was quantified using a Modulation Index (Tort et al., 2010). Power was calculated by filtering for the desired frequency range. For each trial and baseline, the proportion of power in that frequency range to the overall power spectrum was then calculated. Lastly, the trial proportion was then divided by the baseline proportion of the same frequency range.

#### Sharp wave-ripples

Electrophysiology recordings were processed using Neuroexplorer software (Nex Technologies) and custom Python scripts (Laham and Zahn, 2023). For SWR analyses, continuous LFP data were first notched at 60 Hz and band-pass filtered between 140 and 220 Hz. Signal underwent Hilbert transform before being z-scored. Events qualified as SWRs when they exceeded 3 standard deviations for a minimum of 15 milliseconds. SWRs occurring within 15 ms of one another were grouped into a single SWR event. SWR frequency, peak amplitude, integral, duration, and raw number were normalized to each animal’s baseline trial. SWR measures were investigated for baseline, novel, and familiar conspecific trials for the entirety of each testing period.

### Histology

Mice were anesthetized with Euthasol (Virbac) and transcardially perfused with cold 4% paraformaldehyde (PFA). Extracted brains were postfixed for 48 h in 4% PFA at 4◦C followed by an additional 48 h in 30% sucrose at 4◦C for cryoprotection before being frozen in tissue freezing medium at -80◦C. Hippocampal coronal sections (40 μm) were collected using a cryostat (Leica). Sections were pre-blocked for 1.5 hours at room temperature in a PBS solution that contained 0.3% Triton X-100 and 3% normal donkey serum. Sections were then incubated overnight while shaking at 4◦C in the blocking solution that contained combinations of the following primary antibodies: mouse anti-3 microtubule-binding domain tau protein (3R-Tau, 1:500, Millipore, Cat# 05-803), rabbit anti-Purkinje cell protein 4 (PCP4, 1:500, Sigma-Aldrich, Cat# HPA005792), rabbit anti-zinc transporter 3 (ZnT3, 1:500, Alomone labs, Cat# AZT-013), mouse anti SRY-related HMG-box 2 (sox2, 1:100, Millipore, Cat#AB5603), rabbit anti-green fluorescent protein (GFP, 1:500, Molecular Probes, Cat#G10362), or anti-HA tag. For 3R-Tau immunohistochemistry, sections were processed using an antigen retrieval protocol where sections were first incubated in sodium citrate and citric acid buffer for 30 minutes at 80◦C before blocking solution. Washed sections were then incubated for 1.5 hours at room temperature in secondary antibody solutions that contained combinations of the following secondaries: donkey anti-rabbit Alexa Fluor 568 (1:500, Abcam), donkey anti-mouse Alexa Fluor 488 (1:500, Abcam), donkey anti-rabbit Alexa Fluor 488 (1:500, Abcam), or Streptavidin 488 (1:500, Abcam). Washed sections were then counterstained with Hoechst 33342 for 10 minutes (1:5,000 in PBS, Molecular Probes), mounted onto slides, and cover slipped with Vectashield (Vector labs). Slides were coded until completion of the data analysis. Sections through the dorsal hippocampus from electrophysiology studies were stained for Hoechst 33342 to verify accurate and electrode placement.

### Optical intensity measurements

Z-stack images of the CA2 and CA3 were taken using a 40x objective with a 0.5 μm step size through the entire 40 μm thick section on a Leica SP8 confocal with LAS X software (version 35.6). The CA2 was defined by PCP4 labeling, which is specific to the CA2 in the hippocampal CA fields (San Antonio et al., 2014). The CA3 was defined as the region distal to the PCP4 labeling through the hilus. Collected z-stack images were analyzed for optical intensity in Fiji (NIH).

For overall intensity values of 3R-Tau to assess abGC innervation of CA2 and CA3, the ROI included the pyramidal cell layer and stratum radiatum/stratum lucidum. The area, mean gray value, and integrated density of the ROI was calculated for each z-slice and the maximum integrated density value for each z-stack was taken from 3 sections per brain. Averages were obtained for each brain and statistical comparisons were made between groups.

### Cell density/Varicosity density and percentage measurements

The numbers of 3R-Tau+ and sox2+ cells were counted in the dorsal dentate gyrus of the hippocampus on 3 neuroanatomically matched sections using Fiji (ImageJ). The area measurements were determined using Fiji. The densities of 3R-Tau+ cells and sox2+ cells were determined by dividing the total number of positively labeled cells by the volume of the subregion (ROI area multiplied by 40 μm section thickness).

For the GFP+ varicosity analysis, 6 sections per brain were scanned for GFP+ axon segments in the stratum radiatum of the CA2 and CA3. GFP+ varicosities were identified and quantified as circular shaped GFP+ staining that was 0.5–2.2 μm in length and 0.5–1.5 μm in width (Briones et al., 2021, Toni et al., 2008). Proportions of GFP+ varicosities in CA2 compared to CA3 were calculated.

### Statistical Analyses

To investigate potential sex differences in behavioral and brain effects of MSEW, data from males and females were statistically analyzed together with some additional analyses involving individual sexes. For behavioral analyses, discrimination index and total investigation time data sets were analyzed using two-way ANOVAs with sex (male versus female) and rearing condition (MSEW versus control-reared) as variables followed by Tukey post hoc comparisons. Novel and familiar investigation time data sets were analyzed for each sex using two-way repeated measures ANOVAs with Holm-Sidak post hoc comparisons. For histological analyses, 3R-Tau labeled or sox2 labeled cell density, two-way ANOVAs (sex and rearing condition or enriched environment versus standard cage housing as variables) followed by Tukey post hoc comparisons were used. For histological analyses of males alone, data sets were analyzed using unpaired two-tailed Student’s t-tests or Mann-Whitney U tests. For behavioral analyses involving experimental manipulations in adulthood (enriched environment living or chemogenetic activation of abGCs), data sets were analyzed using either a two-way ANOVA or a mixed effects model. Tukey or Holm-Sidak post hoc comparisons were used to follow up any significant main effects or interactions of the ANOVAs. All data sets are expressed as the mean ± SEM on the graphs and statistical significance was set at p < 0.05. GraphPad Prism 9.2.0 (GraphPad Software) and R studio were used for statistical analyses. All graphs were prepared using GraphPad Prism 9.2.0 (GraphPad Software).

## Results

### MSEW impairs social recognition in male mice and improves social novelty preference in female mice

Using the direct social interaction test, we examined social recognition in control-reared and MSEW adult male and female mice (Figure 1A). Because control-reared mice investigate novel mice more than familiar mice, social recognition was defined as a significant decrease in investigation time between novel and familiar trials. Analyses of the discrimination indices (novel minus familiar divided by novel plus familiar) revealed a significant interaction between sex and rearing condition with MSEW male but not female mice showing an impairment in social recognition (F (1, 47) = 6.978; p=0.0112) but no main effect of sex or rearing condition. Post hoc tests showed that MSEW males had lower discrimination indices than control-reared males (p= 0.0086) while MSEW females were not statistically different from control-reared females (p=0.9626) (Figure 1B). Analyses of the overall investigation times showed that there was a sex by rearing condition interaction (F (1, 46) = 4.812; p=0.0334), a main effect of sex (F (1, 46) = 32.430; p<0.0001) but not rearing condition (F (1, 46) = 0.1076; p=0.7444). Post hoc comparisons revealed no significant difference between control and MSEW males (p= 0.2871) nor between control and MSEW females (p= 0.1996) (Figure 1C). Taken together, these findings suggest that MSEW impairs social recognition in adult males and that this effect is not driven by a change in overall investigation times. Analyses of individual sex data showed an interaction between rearing condition and social stimuli (F (1, 27) = 9.274; p=0.0051) with control-reared male mice showing significantly lower investigation times for the familiar mouse than the novel mouse (p<0.0001), while MSEW male mice did not have significant differences in interaction time across trials (p=0.1270) (Figure 1D). Analyses of female behavioral data also showed a significant interaction between rearing condition and social stimuli (F (1, 19) = 6.292; p=0.0214) but both female control-reared and MSEW mice had significant declines in investigation time between novel and familiar trials (control-reared, p=0.0280; MSEW, p<0.0001). (Figure 1E). Also, in contrast to what was observed with male mice, MSEW females showed significantly higher investigation of novel mice compared to control-reared females (p=0.0009) (Figure 1E).

**Figure 1:**
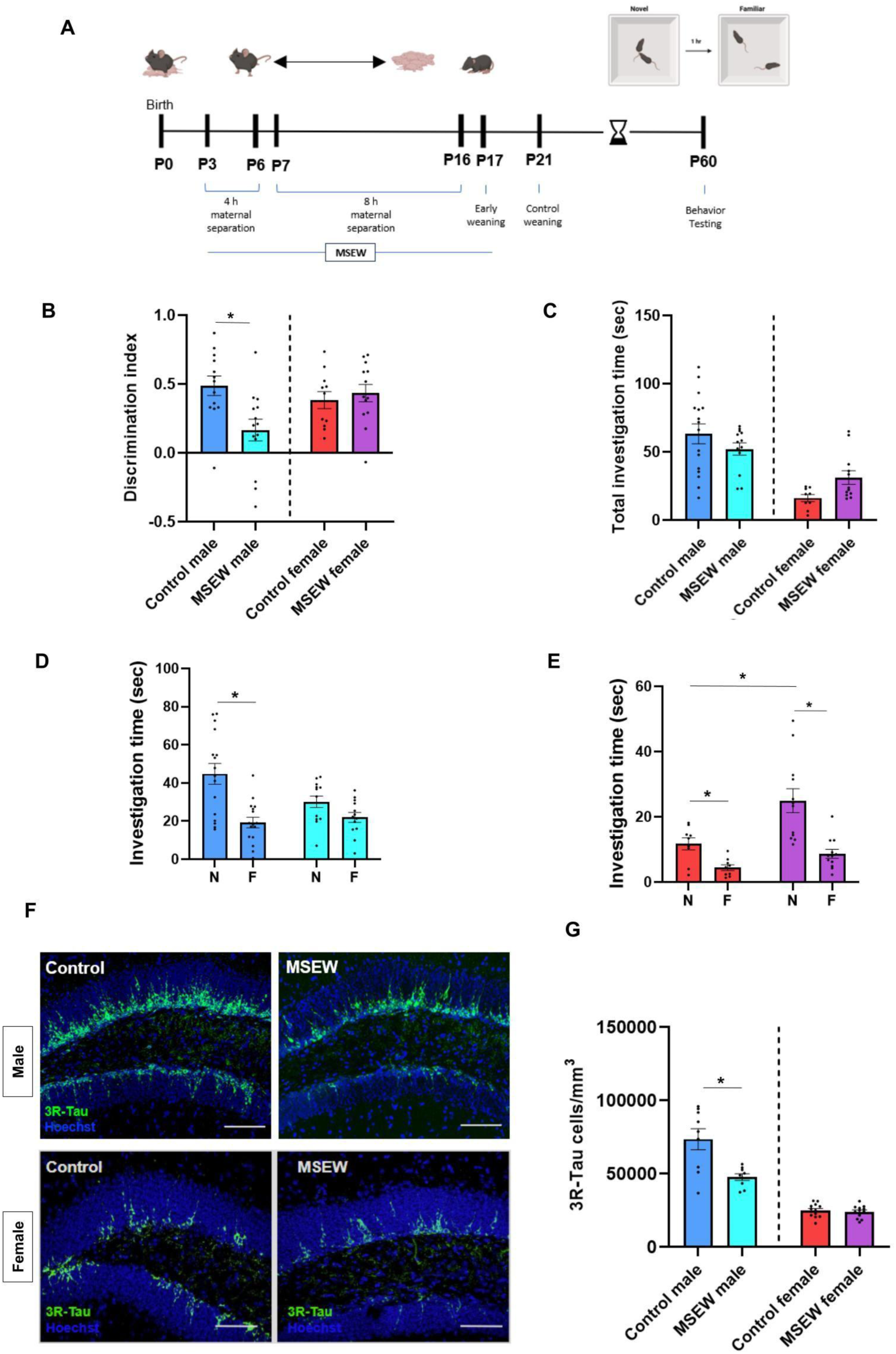
MSEW impairs social recognition and adult neurogenesis in male mice. **A**) Experimental timeline and schematic of MSEW and social memory testing. **B**) MSEW males had lower discrimination indices (N-F/N+F) than control-reared males while MSEW females were not significantly different on this measure from control-reared females. **C**) Total investigation time (N+F) is not different between control-reared and MSEW males nor between control-reared and MSEW females. **D**) Control-reared males display significant reductions in time between investigating the novel mouse in trial 1 and the familiar mouse in trial 2 but MSEW males do not show this reduction. **E**) Control-reared and MSEW females both show a significant reduction in investigation time between novel and familiar stimulus mouse exposure with an increase in investigation time of novel mice observed in MSEW females. **F**) Confocal images of 3R-Tau labeled immature neurons in the dentate gyrus of males and females showing fewer labeled cells in MSEW males compared to control males but no differences between MSEW females and control females. Scale bars = 50 μm. **G**) Adult male MSEW mice have fewer 3R-Tau labeled immature neurons in the dentate gyrus than control males with no changes in this measure detected in MSEW females compared to control females. *p<0.05, n=11-16 for male behavioral studies; n=8-13 for 3R-Tau measurements. Abbreviations: maternal separation with early weaning (MSEW), postnatal day (P), novel (N), familiar (F)

### MSEW results in fewer immature neurons in the dorsal dentate gyrus of adult males but not adult females

Evidence suggests that abGCs are critical for social recognition in healthy mice (Cope et al., 2020) and their afferents to the CA2 have been shown to facilitate social memory retrieval (Laham et al., 2024). Whether MSEW affects adult-born granule cells, their progenitor cells, and their projections to CA2 in males and females was investigated. Using 3R-Tau to label immature granule cells in dorsal dentate gyrus (Llorens-Martin et al., 2015), we found a significant interaction between sex and rearing condition (F (1, 39) = 14.19; p=0.0005) with significant main effects of sex (F (1, 39) = 119.0, p<0.0001) and rearing condition (F (1, 39) = 16.43, p=0.0002). Post hoc comparisons revealed that MSEW males had significantly fewer 3R-Tau+ cells than control-reared males (p <0.0001) while MSEW females did not show any difference from control-reared females in the number of 3R-Tau+ abGCs (p= 0.9961) (Figure 1F,G).

Because we observed a significant decrease in immature neurons in MSEW males compared to control-reared males, we explored this difference further by quantifying the intensity of 3R-Tau axonal labeling in two target areas of dentate gyrus granule cells, the stratum radiatum of the CA2 region and the stratum lucidum of the CA3 region. Consistent with the finding of fewer 3R-Tau labeled cells in the MSEW dentate gyrus, we found reduced 3R-Tau intensity in both CA2 (t13=3.090; p=0.0086) and CA3 (t15=2.481; p=0.0254) (Figure S1A-D).

Because adult male MSEW mice had lower numbers of immature neurons in the dentate gyrus and correspondingly diminished axons in two target regions, we then investigated whether adult MSEW males had fewer progenitor cells in the dentate gyrus. Using sox2 as a marker, we found that MSEW males had fewer progenitor cells in the dentate gyrus than control-reared males (t15=2.460; p=0.0265) (Figure S1E,F). We next labeled adult-born granule cells in MSEW and control-reared males with a GFP-retrovirus and investigated whether axonal varicosities, sites of en passant synapses, were differentially affected by MSEW in these target regions. We found that while CA2 and CA3 have roughly similar densities of these putative synaptic contacts in control-reared males, this proportion shifted in MSEW males, such that a greater loss was detected in CA2 (t6=3.546; p=0.0121) (Figure S1G,H).

Next, to assess whether the MSEW-induced reduction in dentate gyrus projections to the CA2 and CA3 is specific to immature abGC axons or whether a similar reduction would exist in mature granule cell axons, which include developmentally born cells, the zinc transporter ZnT3 was used to label mature granule cell axons. Control-reared and MSEW males showed no difference in intensity of ZnT3+ axons in the stratum radiatum of the CA2 (t16=0.4873, p=0.6326) or in the stratum lucidum of CA3 (t14=1.653, p=0.1207) (Figure S1I-K). Taken together, these findings suggest that in adult males, MSEW is associated with diminished afferents from adult-born immature neurons to both CA2 and CA3, but not an overall reduction in mature granule cell afferents to these areas.

### Stimulating adult neurogenesis with enriched environment living restores social recognition in MSEW male mice

Previous studies have shown that living in an environmental enrichment stimulates adult neurogenesis in the dentate gyrus of mice and rats (Olson et al., 2006; Speisman et al., 2013; Monteiro et al., 2014). Given that we found lower numbers of abGCs and impaired social recognition in MSEW males, we hypothesized that living in an enriched environment might boost adult neurogenesis and thereby improve social recognition. In a new cohort of mice, males that underwent MSEW were moved to enriched environment housing or remained in standard housing in young adulthood (Figure 2A). Six weeks later, mice were tested in the direct social interaction test. MSEW mice living in standard housing again showed impaired social recognition when investigating a sex- and age-matched stimulus mouse, while standard housing control-reared mice investigated a novel mouse significantly more. However, this impairment was rescued in the MSEW males by enriched environment housing with no effect on control-reared mice housed in environmental enrichment (Figure 2B). A significant interaction was observed between rearing group/housing group and social stimuli (F (3, 43) = 6.129; p=0.0015) with significant post hoc comparisons showing social recognition in the MSEW group with enriched environment living (p <0.0001) but not standard housing (p= 0.9520). Analyses of discrimination ratios also followed this pattern; there was a significant interaction between rearing condition and adult housing condition (F (1, 43) = 16.21; p=0.0002) with significant main effects of both rearing (F (1, 43) = 4.239, p=0.0456) and housing (F(1,43) =13.87; p=0.0006). Post hoc comparisons showed significantly lower discrimination indices in the MSEW standard housing condition compared to MSEW enriched environment condition (p<0.0001) (Figure 2C). Two-way ANOVAs revealed significant interactions between rearing condition and adult housing condition for sox2+ progenitor cells (F (1, 37) = 4.303; p=0.0451) (Figure 2D,E) and 3R-Tau+ immature granule cells (F (1, 41) = 4.081; p=0.0499) (Figure 2F,G) in the dentate gyrus. Enriched environment living increased the number of sox2+ cells and 3R-Tau+ cells in the dentate gyrus of MSEW male mice so that they were significantly higher than MSEW mice living in control conditions (sox2: p = 0.0089; 3R-Tau: p= 0.0135) and not different from control-reared groups (Figure 2D-G). These findings suggest that an experience-dependent increase in the number of progenitor cells and abGCs of the dentate gyrus may be sufficient to restore social recognition in MSEW males.

**Figure 2:**
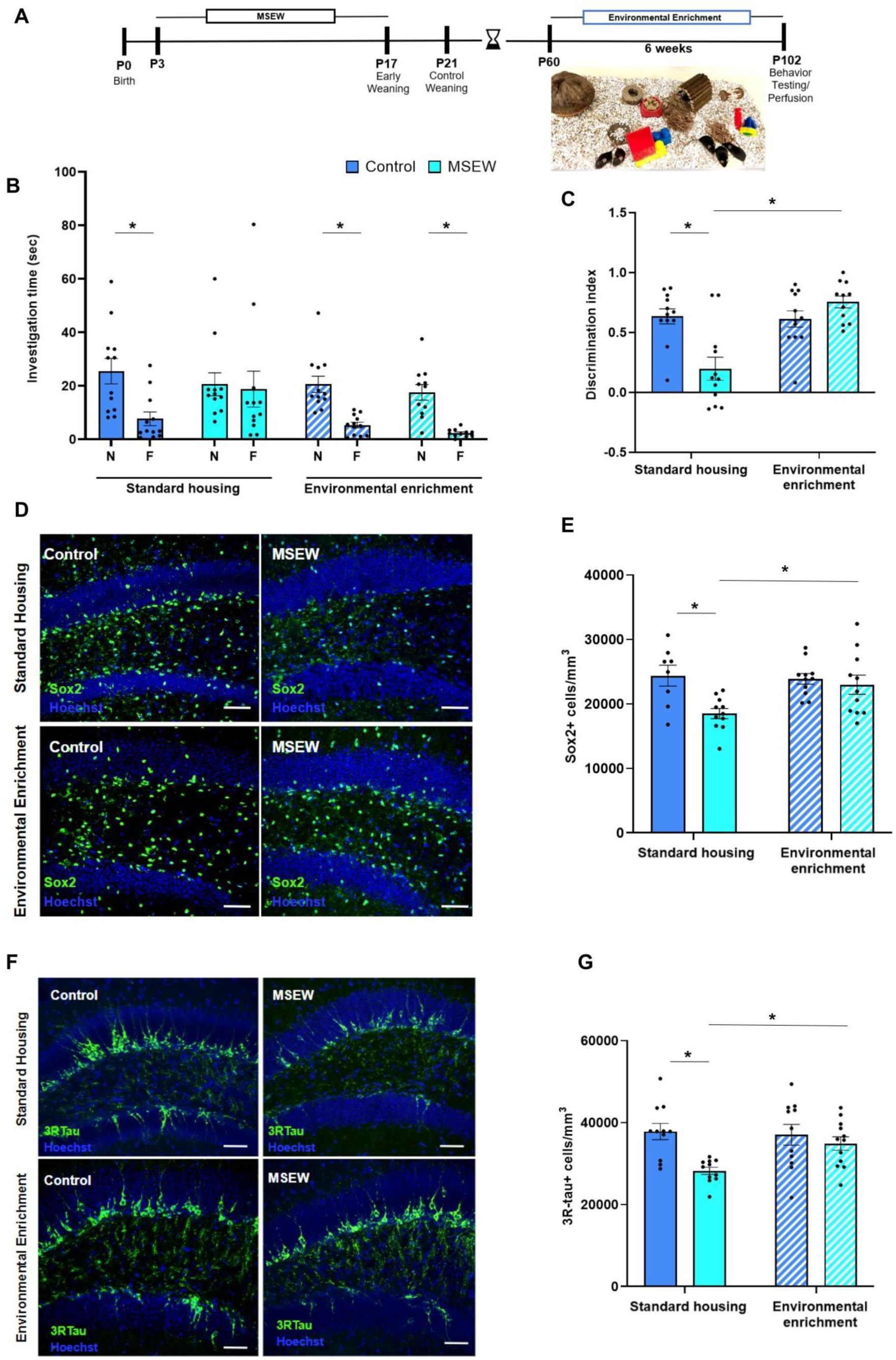
Enriched environment housing in adulthood restores MSEW-induced effects on social recognition, hippocampal progenitor cells, and adult-born granule neurons. **A)** Timeline of MSEW/enriched environment experiment including image of enriched environment housing. **B)** Control-reared/standard housing adult males display a significant reduction in time investigating the novel mouse in trial 1 to the familiar mouse in trial 2. MSEW/ standard housing adult males had similar investigation times in trial 1 and trial 2. Environmental enrichment control and MSEW males display a significant reduction in time investigating the novel animal in trial 1 to the familiar animal in trial 2. **C)** MSEW/standard housing males had a significantly lower discrimination index compared to control-reared/standard housing males and MSEW/environmental enrichment males. **D**) Confocal images of the dorsal dentate gyrus showing examples of sox2 labeling in all 4 treatment groups. Scale bars = 30 μm. **E)** MSEW/standard housing males had significantly lower sox2+ cell density in the dorsal dentate gyrus compared to control-reared/ standard housing, control-reared/environmental enrichment and MSEW/environmental enrichment males. **F**) Confocal images of the dorsal dentate gyrus showing examples of 3R-Tau labeling in all 4 treatment groups. Scale bars = 30 μm. **G**) MSEW/standard housing males had significantly fewer 3R-Tau+ cells in dorsal dentate gyrus compared to control-reared/standard housing males, control-reared/environmental enrichment males, and MSEW/environmental enrichment males. *p<0.05. n=12 for behavioral studies; n=8-12 for sox2 and 3R-Tau measurements. Abbreviations: maternal separation with early weaning (MSEW), postnatal day (P), novel (N), familiar (F)

### Chemogenetic activation of abGCs restores social recognition in MSEW male mice

To more specifically investigate the role of abGCs in social recognition after MSEW, tamoxifen-inducible nestin-Cre double transgenic mice with abGCs transduced to express excitatory Gq DREADDs were generated. This allowed for the temporary activation of these abGCs at different points after their generation by injecting the ligand CNO. Previous studies have shown that abGCs exert an important influence over hippocampal microcircuitry between 4-6 weeks after their generation, with their influence diminishing after this time (Laham et al., 2024). To test whether activating abGCs of different ages after their generation mitigates social recognition dysfunction after MSEW, adult male control-reared and MSEW single transgenic (ST, nestin-Cre) and double transgenic (DT, nestin-Cre:Gq) mice were treated with tamoxifen and tested on the direct social interaction task 4-6 weeks after abGC labeling (Figure 3A,B). Thirty minutes before each behavioral trial, mice were injected with either vehicle or CNO. Each mouse was tested twice, once with vehicle and once with CNO, with injections counterbalanced across tests with a 48-hour washout period between tests. Examination of trial-by-trial behavioral data using a repeated measures design in vehicle-treated mice showed a significant interaction between rearing condition and social recognition behavior (F (3, 51) = 14.56; p<0.0001) with both ST and DT MSEW groups showing no significant difference (ST: p= 0.2898; DT: p= 0.3079) while both control-reared groups showed significant differences (ST: p<0.0001; DT: p<0.0001) between novel and familiar social investigation (Figure 3C). Treatment with CNO also showed a significant interaction between rearing condition and social recognition (F(3,48) = 8.268; p=0.0002) but with significant differences between novel and familiar social investigation observed in both control-reared groups (ST: p <0.0001; DT: p<0.0001) as well as the MSEW DT group (p<0.0001). MSEW ST mice remained unchanged by CNO treatment with no significant difference between novel and familiar investigation times (p= 0.9767) (Figure 3D). Analysis of discrimination indices confirmed these findings with a significant interaction between transgenic line and rearing condition/CNO administration (F (3, 105) = 9.723; p<0.0001) with significant differences between control-reared and MSEW mice in ST and DT mice treated with vehicle treatment (ST:p= 0.0002; DT: p=0.0497), as well as in ST mice with CNO treatment (p<0.0001) (Figure 3E). By contrast, treatment of MSEW DT mice with CNO increased the discrimination index such that no significant difference was observed when compared to control-reared DT mice (p= 0.9527). Accordingly, MSEW DT mice treated with CNO were significantly different from MSEW DT mice treated with vehicle (p= 0.0027) (Figure 3E).

**Figure 3:**
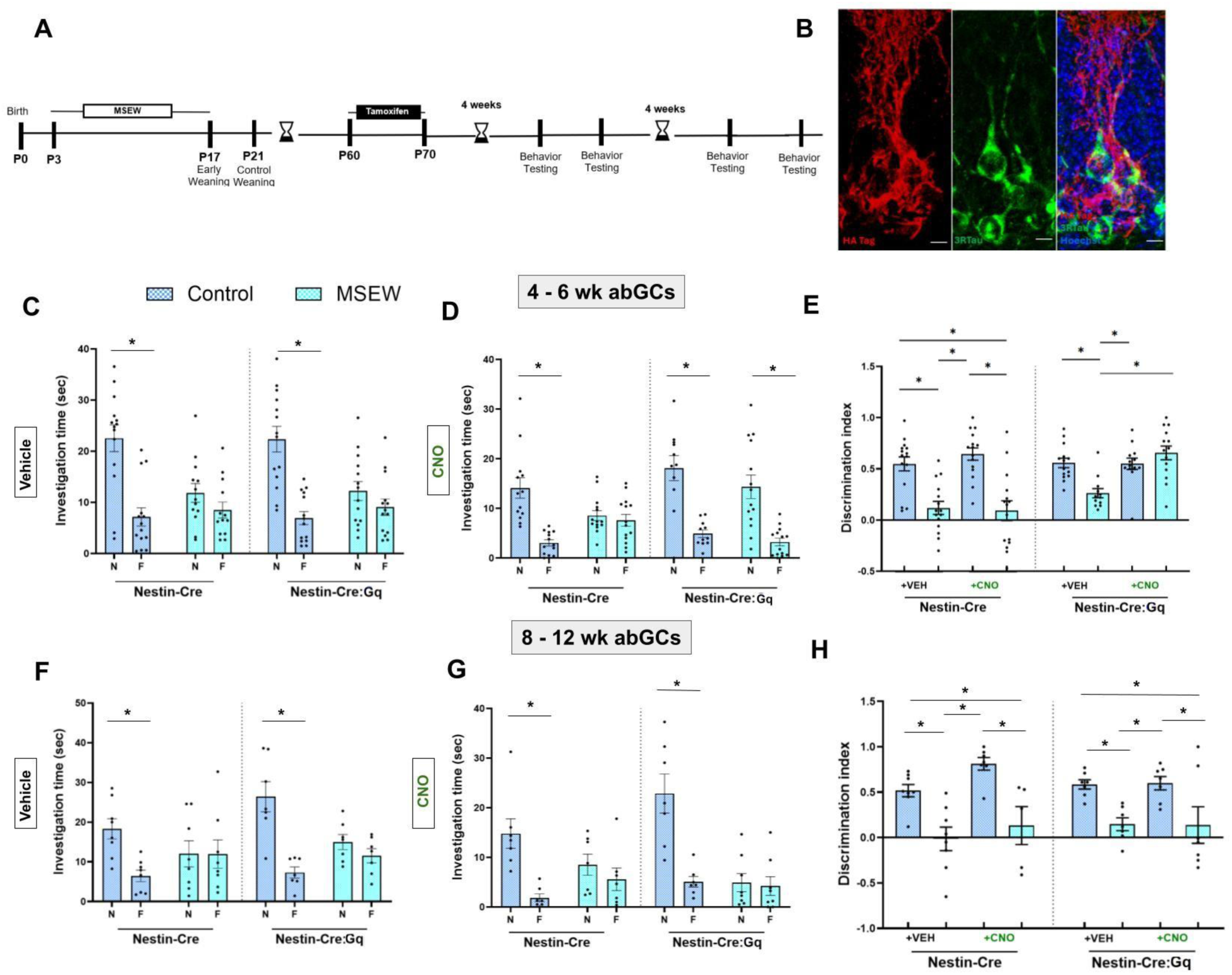
Activation of 4-6 week old, but not 8-12 week old, abGCs restores social recognition in MSEW males. **A)** Timeline of MSEW double transgenic experiment. **B)** Confocal images of HA-tag labeled neurons co-labeled with the immature neuron marker 3R-tau in the dentate gyrus of an adult male nestin-Cre:Gq mouse. Scale bars = 10 μm. **C**) After vehicle injection, control-reared nestin-Cre and nestin-Cre:Gq with a vehicle injection both showed a significant reduction in time investigating the novel mouse in trial 1 to the familiar mouse in trial 2. MSEW nestin-Cre and nestin-Cre:Gq both showed similar investigation times in trial 1 and trial 2. **D)** After CNO injection, control-reared nestin-Cre, control-reared/nestin-Cre:Gq, and MSEW/ nestin-Cre:Gq mice all showed a significant reduction in time investigating the novel animal in trial 1 (N) to the familiar animal in trial 2 (F). MSEW nestin-Cre continued to show similar investigation times in trial 1 and trial 2, with CNO injection. **E)** With a vehicle injection, MSEW nestin-Cre and nestin-Cre:Gq both had significantly lower discrimination indices compared to control-reared nestin-Cre and nestin-Cre:Gq. However, after CNO injections, only MSEW nestin-Cre mice had significantly lower discrimination indices than control-reared nestin-Cre, control-reared nestin-Cre:Gq, and MSEW nestin-Cre:Gq mice. **F)** Activating 8-12 week old abGCs does not rescue social recognition. Control-reared nestin-Cre and nestin-Cre:Gq with a vehicle injection both showed a significant reduction in time investigating the novel animal in trial 1 to the familiar animal in trial 2. MSEW nestin-Cre and nestin-Cre:Gq both showed similar investigation times in trial 1 and trial 2, with vehicle injection. **G)** Control-reared nestin-Cre and nestin-Cre:Gq with a CNO injection both showed a significant reduction in time investigating the novel animal in trial 1 to the familiar animal in trial 2. MSEW nestin-Cre and nestin-Cre:Gq both showed similar investigation times in trial 1 and trial 2, with vehicle injection. **H)** With a vehicle injection, MSEW nestin-Cre and nestin-Cre:Gq both had significantly lower discrimination indices compared to control-reared nestin-Cre and nestin-Cre:Gq. Activating 8-12 week old abGCs with CNO had no effect on either MSEW or control-reared mice. *p<0.05. n=14-15 for 4-6 week abGC behavioral studies; n=7-8 for 8-12 week abGC behavioral studies. Abbreviations: maternal separation with early weaning (MSEW), postnatal day (P), novel (N), familiar (F), adult-born granule cells (abGCs).

The beneficial effects of CNO on social recognition in MSEW DT mice were no longer observed when abGCs were 8-12 weeks of age (Figure 3F-H). In this experiment, the difference in social recognition between control-reared and MSEW ST and DT mice remained the same regardless of whether mice were treated with CNO or not (vehicle ST: p= 0.0230, vehicle DT: p= 0.0096; CNO ST: p= 0.0094; CNO DT: p=0.0042) with no difference between MSEW DT treated with vehicle versus CNO (p>0.9999). Collectively, these findings suggest that activation of abGCs during a certain time window after their generation (4-6 weeks) but not later is sufficient to restore social recognition deficits in MSEW male mice.

### CA2 network activity is altered by MSEW and chemogenetic activation of abGCs

Studies have shown that CA2 network activity is associated with social recognition (Oliva et al., 2020; Zhu et al., 2023) and activation of dentate gyrus abGCs alters CA2 network activity (Laham et al., 2024). Whether MSEW affects CA2 network activity and whether any such effects would be restored by chemogenetic activation of abGCs was investigated next. For this experiment, control-reared and MSEW double transgenic males implanted with electrodes in the unilateral CA2 were examined under conditions of CNO or vehicle injection 4-6 weeks after tamoxifen injection to express excitatory DREADDs in abGCs. First, we examined whether changes in theta (4-12 Hz) or gamma (25-130 Hz) power differed among control-reared and MSEW mice with vehicle or CNO treatment. We found no significant interactions between rearing condition and drug treatment on any of these measures during exposure to the novel stimulus mouse or the familiar stimulus mouse (Figure S2A-H). Because phase amplitude coupling (PAC) between theta and gamma has been associated with social recognition (Zhu et al., 2023; Laham et al., 2024), we examined PAC between theta and slow (25-50 Hz), mid (50-100 Hz), or fast (100-130 Hz) gamma. We found no significant interactions between rearing condition and drug treatment on any of these measures during exposure to the novel stimulus mouse or the familiar stimulus mouse (Figure S3A,B). These findings suggest that despite differences in social recognition between control-reared and MSEW males, there were no detectable differences in theta/gamma power or PAC with MSEW with either vehicle or CNO treatment.

CA2 SWRs have been linked to social recognition (Oliva et al., 2020) so we next investigated whether they differed between control-reared and MSEW male mice with and without CNO (Figure 4A-C). We found a significant interaction between rearing condition and drug with SWR rate (F (1, 7) = 12.03; p=0.0104). Post hoc comparisons showed that SWR rate was lower in MSEW males treated with vehicle compared to control-reared males (p= 0.0047) with no significant difference between rearing condition after administration of CNO (p=0.2225) (Figure 4D). Like previously reported findings (Laham et al., 2024), when treated with vehicle, control-reared mice had higher CA2 SWR rate while investigating novel mice compared to familiar mice analyzed using discrimination indices (Figure 4E). This difference did not exist with MSEW male mice treated with vehicle but was restored with CNO treatment. Analyses of these data also showed a significant interaction between rearing condition and drug (F (1, 6) = 7.504; p=0.0338) with significantly lower discrimination index in the MSEW vehicle group compared to control-reared vehicle group (p=0.0025) and a significant restoration observed in MSEW mice after CNO treatment (p=0.0078) (Figure 4E). MSEW appeared to prevent the differential SWR rate between novel and familiar stimulus exposure, an effect that was reversed by activation of abGCs with CNO. We next examined other properties of SWRs across groups, including peak amplitude, duration, and integral. We found no significant interactions between rearing condition and DREADD activation in SWR amplitude during any recording condition (Figure S4A,B), but several significant interactions were observed in SWR duration and integral during social exposures. During novel and familiar mouse exposure, significant interactions were observed in SWR duration (novel: F (1, 7) = 11.67; p=0.0112; familiar: F (1, 7) = 11.20; p=0.0123). During both types of exposure, SWR duration was longer in MSEW than control-reared vehicle-treated mice (novel: p=0.0037; familiar: p=0.0019) but with no significant change following CNO treatment. SWR duration was increased by CNO treatment but only in the control-reared group (novel: p=0.0080; familiar: p=0.0397) (Figure 4F,G). During novel mouse exposure, a significant interaction between rearing condition and DREADD activation was observed in the SWR integral (F (1, 7) = 8.252; p=0.0239). SWR integral was elevated in MSEW compared to control-reared mice (p= 0.0303) but CNO treatment only changed the SWR integral in control-reared mice (p= 0.0088) (Figure 4H). Because the SWR integral is a metric that combines the amplitude and duration of SWRs, it is likely that these latter differences reflect changes in duration, but not amplitude, of SWRs. Taken together, these findings suggest that MSEW reduces SWR rate during novel social encounters, as well as the difference in SWR rate between novel and familiar exposures, and that the difference in SWR rate is corrected by DREADD activation. SWR duration and integral are greater in MSEW than control-reared mice but these measures are not corrected in MSEW mice by DREADD activation.

**Figure 4:**
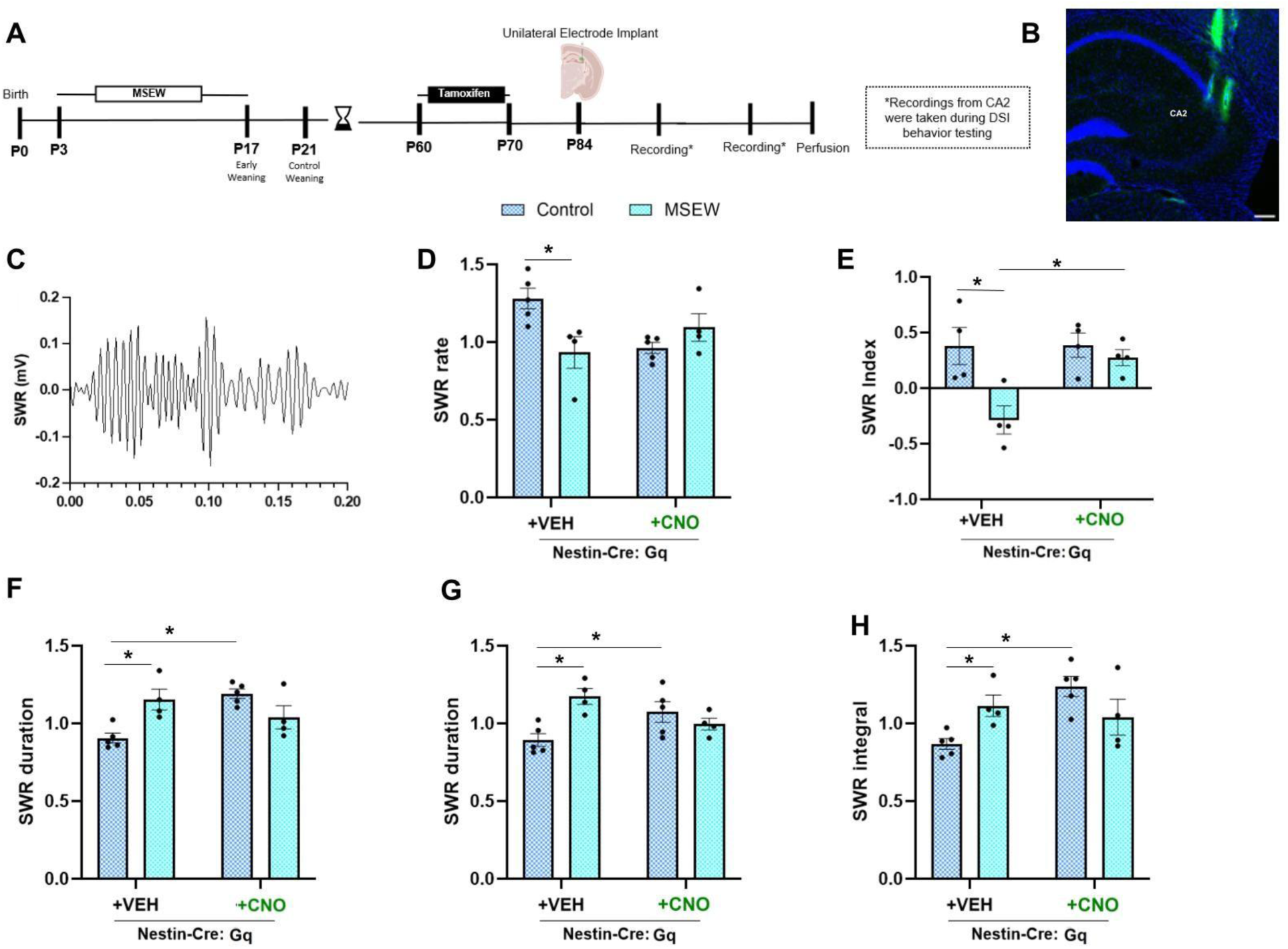
Activation of 4-6 week old adult-born granule cells restores CA2 network activity in MSEW males. **A)** Timeline of MSEW chemogenetic electrophysiology experiment. **B)** Confocal image of representative electrode implant in CA2 with electrode track in green. Scale bar = 20 μm. **C)** Filtered SWR example trace from one mouse. **D)** With vehicle injection, MSEW nestin-Cre:Gq animals had reduced SWR rate during exposure to the novel mouse (Trial 1). This reduction was not significant after CNO injection. **E)** With a vehicle injection, MSEW nestin-Cre:Gq males had a significantly lower SWR index than VEH: Control nestin-Cre:Gq. After CNO injection, this difference was eliminated. **F)** With a vehicle injection during trial 1, MSEW nestin-Cre:Gq had longer SWR duration than control-reared nestin-Cre:Gq mice. This difference was not eliminated with CNO injection. **G)** With a vehicle injection during trial 2, MSEW nestin-Cre:Gq had longer SWR duration than control-reared nestin-Cre:Gq mice. This difference was not eliminated with CNO injection. **H**) With a vehicle injection, MSEW nestin-Cre:Gq had greater SWR integral during trial 1 than control-reared nestin-cre:Gq mice. This difference was not eliminated with CNO injection. *p<0.05. n=4-5 mice (4 electrodes per mouse). Abbreviations: maternal separation with early weaning (MSEW), postnatal day (P), novel (N); familiar (F), sharp wave ripple (SWR).

## Discussion

We found that MSEW impaired social recognition in adult males but not females, which instead showed an increase in social novelty investigation. Along with impaired social recognition, compared to control-reared mice, MSEW males, but not females, had fewer abGCs in the dentate gyrus. In addition, MSEW males had fewer progenitor cells in the dentate gyrus, as well as fewer abGC axons in both the CA2 and CA3 regions. We also found altered CA2 network activity in MSEW males compared to control-reared males, including reduced SWR rate, smaller differential SWR rate in response to novel compared to familiar social stimuli, and greater SWR duration. Additionally, enriched environment living restored the number of progenitor cells and abGCs in the dentate gyrus of MSEW males to control male levels, which was accompanied by an improvement in social recognition. Chemogenetic activation of abGCs in MSEW males also restored social recognition and this beneficial effect was accompanied by a normalization of the differential CA2 SWR rate in response to social stimuli. Taken together, these findings suggest that adult interventions to enhance abGC number or activity in MSEW males are sufficient to restore behavioral and CA2 network function.

### MSEW impairs social recognition through suppressed adult neurogenesis in the dentate gyrus

Previous studies have shown that abGCs in the dentate gyrus are important for social recognition in mice. The ability of adult mice to recognize a mouse as familiar 24 hours after the first encounter is impaired in mice lacking abGCs (Cope et al., 2020). Similar impairments in the ability to discriminate between a familiar mouse encountered during development (i.e., the mother) and a novel mouse have been observed in mice with ablated abGCs (Laham et al., 2024). These findings are consistent with our observations that MSEW male mice have both impaired social recognition and reduced numbers of abGCs. In addition, our results showing that enriched environment living restores numbers of abGCs in MSEW mice to control levels along with recovery of social recognition function are consistent with the hypothesis that reduced numbers of abGCs are causally linked to MSEW-induced social dysfunction. Because enriched environment living stimulates multiple modes of plasticity in addition to adult neurogenesis (van Praag et al., 2000; Hosseiny et al., 2015; Foggetti et al., 2019), we used a double transgenic model to investigate whether activating abGCs, without altering their density, would also restore social dysfunction after MSEW. Like our behavioral results with enriched environment housing, activating abGCs in MSEW males restored social recognition. Taken together, these findings strongly suggest that MSEW deficits in social recognition are causally linked to reduced abGCs, effects that can be reversed in adulthood by restoring the number, or increasing the activation, of abGCs.

Studies have shown that in addition to their well-established connections to the hilus and the CA3 region, abGCs project to the CA2 region (Llorens-Martin et al., 2015), an area known to be important for social recognition (Hitti and Siegelbaum, 2014). A previous study investigating this projection using an inhibitory DREADD model, including CNO delivery to CA2, showed the importance of this circuit in discriminating mothers from novel mothers in adult offspring mice (Laham et al., 2024). The current study enhanced abGC activity generally, by giving CNO systemically, so we cannot rule out the possibility that abGC projections to the CA3 are involved in the restoration of MSEW-induced deficits in social recognition. Indeed, we found reduced abGC axons in the CA2 and CA3 of MSEW males, although a greater reduction in varicosities was observed in CA2 than CA3. It is possible that the combined effect of abGCs on social recognition involve both projections to CA2 and CA3, as well as communication between these regions.

Our data further suggest that the reduction in number of abGCs and their axons in target regions in MSEW males is likely due to a deficit in cell proliferation, as opposed to cell survival, as MSEW males have fewer sox2+ progenitor cells in the dentate gyrus than control-reared mice, an effect that is normalized by enriched environment living. The restoration of sox2+ cell numbers with enriched environment living is likely caused by increased proliferation of progenitor cells, whose expansion would ultimately increase the number of abGCs. These findings are partially consistent with previous reports showing that enriched environment living stimulates cell proliferation and the number of sox2+ cells in the dentate gyrus, although these previous studies showed this effect in control rodents (Kempermann et al., 1997; Veena et al., 2009; Tanti et al., 2012; Chandler et al., 2020) and we only observed the increase in MSEW mice, possibly due to differences in species, strain, sex, or age of the experimental animals, as well as differences in features of the enriched environment .

Since MSEW is a developmental experience that is completed long before adulthood, our findings raise questions about how the hippocampus might be affected during the adverse experience to ultimately have fewer progenitor cells available to generate new neurons in adulthood. One possibility is that MSEW accelerates cell proliferation and neurogenesis such that the progenitor pool is depleted sooner, findings that are consistent with some developmental stress postnatal neurogenesis studies (Naninck et al., 2015). Accelerated cell proliferation may reflect an advanced developmental aging process that has been reported for some early life adversity models (Bath et al., 2016; Chaudhari et al., 2022). For example, studies have shown that age-related decline in adult neurogenesis is causally linked to the expression of senescence molecules by progenitor cells in the hippocampus (Fatt et al., 2022). Taken together with the early life stress literature on accelerated aging, it is possible that the MSEW male dentate gyrus in young adulthood is more like a control-reared male at an older age. Importantly, MSEW effects can be reversed by housing mice in enriched environments, which has also been observed with other age-related phenomenon in control-reared mice (Birch and Kelly, 2019; Singhal et al., 2019; Schmidt et al., 2022). It has been hypothesized that early life adversity-induced accelerated aging could be adaptive if it hastens maturity and independence, although several studies have shown that early life adversity slows the achievement of developmental milestones, including puberty (Knop et al., 2019; Manzano Nieves et al., 2019).

### CA2 network activity and social recognition

A growing number of studies have shown that CA2 network activity is associated with healthy social recognition capabilities. SWRs and phase amplitude coupling in the CA2 have been linked to social experience, social memory consolidation and retrieval (Oliva et al., 2020; Zhu et al., 2023; Laham et al., 2024; Diethorn et al., 2026). CA2 SWRs are known to be causally linked with social memory consolidation through their coordination with the replay of neuronal ensembles (Oliva et al., 2020). In the CA2, SWRs are more likely to occur during quiet wakefulness than during sleep compared to other hippocampal subregions and are elevated during exposure to a novel versus familiar social stimulus (Laham et al., 2024). In control-reared males treated with vehicle we found a similar pattern of differential CA2 SWR responses depending on whether the social stimulus was novel or familiar. Nestin-Cre:Gq males previously subjected to MSEW and treated with vehicle lacked the differential SWR response to novel compared to familiar social stimuli. When nestin-Cre:Gq MSEW mice were treated with CNO to activate abGCs, the novel/familiar difference in SWRs was evident, such that no differences were observed between nestin-Cre:Gq control-reared and MSEW males. Adult-generated neurons in the dentate gyrus play a significant role in social cognition, including facilitating sparse encoding and participating in the generation of SWRs in the CA2 (McHugh et al., 2022; Laham et al., 2024). A previous study showed that inhibiting the abGCs-CA2 pathway in control-reared mice leads to decreased CA2 SWR diversification (Laham et al., 2024), which is the same CA2 SWR pattern we observed with MSEW. Taken together, MSEW-induced decreases in abGCs may impair social memory through homogenous SWRs during encoding and retrieval of social memories. MSEW also changed CA2 SWR characteristics, including duration and integral, but neither of these differences were corrected by activation of abGCs, suggesting that some features of CA2 network activity may be more closely coupled to healthy social recognition than others.

### Sex differences in MSEW effects

Our results demonstrate sex differences in behavioral response to MSEW. In adulthood, MSEW males have impaired social recognition. In striking contrast, adult female mice exhibit robust social recognition abilities, with females showing higher social novelty responses compared to control-reared females. The behavioral effect of MSEW on females is likely not due to changes in abGCs, suggesting a different unknown mechanism underlies this effect. These findings raise questions about whether the sex differences arise from differential developmental experience or whether they emerge over time. Regarding the former possibility, a few studies have shown that rodent mothers display differential care toward male and female pups, perhaps due to a greater need for licking and grooming to stimulate micturition in males (Maggi et al., 1986; Sakamoto et al., 2021). Thus, maternal separation, which alters maternal behavior (Bailoo et al., 2014; Baracz et al., 2020) may do so differently toward male and female pups, with greater long-term effects observed in males. However, recent evidence suggests that the effects of MSEW are largely similar on male and female pups, but that females recover as time passes, possibly due to influences that occur during puberty (Bapat et al., 2026). Understanding mechanisms that promote recovery from ELA-induced social recognition impairment in females may suggest additional points of intervention for recovery in MSEW males.

### Potential translational relevance

Our findings suggest that behavioral outcomes after MSEW are influenced by sex. Similar sex-specific behavioral effects were observed with the limited bedding and nesting (LBN) manipulation, a developmental model of scarcity, with social recognition impairments in adult males but not females (Waters et al., 2026) Collectively, these findings suggest that early life adversity may exert a general effect on social memory circuitry, but available evidence suggests that these similar behavioral outcomes are supported by different brain changes. The current study identified reduced numbers of abGCs, with their connections to the CA2, as a cause of MSEW-induced social recognition impairment in males. The LBN manipulation, however, did not produce changes in abGCs in males despite deficits in social recognition. Instead, LBN produced an increase in perineuronal nets, specialized extracellular matrix structures that inhibit plasticity, in the CA2 region, which when reduced through targeted intervention in adulthood, restore social memory. Thus, two different types of early adverse experiences affect the same behavior by engaging separate plasticity mechanisms in overlapping brain regions. These findings are consistent with some human literature showing that different types of childhood maltreatment predispose individuals to develop certain types of neuropsychiatric illnesses over others, and that different neural circuits are altered as well (Waters and Gould, 2022). While childhood experience in humans is undeniably more complex than early-life adversity manipulations in laboratory rodents, our findings suggest that early life stress is not a general experience that results in similar brain changes regardless of individual experience. Understanding how different early-life adversity paradigms alter the brain during development using rodent models may yield potential biomarkers, and therapeutic targets for reversing dysfunction prior to adulthood and may help in the development of precision medicine approaches for assisting individuals with neuropsychiatric disease after childhood adversity.

## Acknowledgments

This work was supported by the National Institute of Mental Health R01MH117459 to E.G. and National Science Foundation Graduate Research Fellowship Program 2021318039 to R.C.W.

---

**Figure S1:**
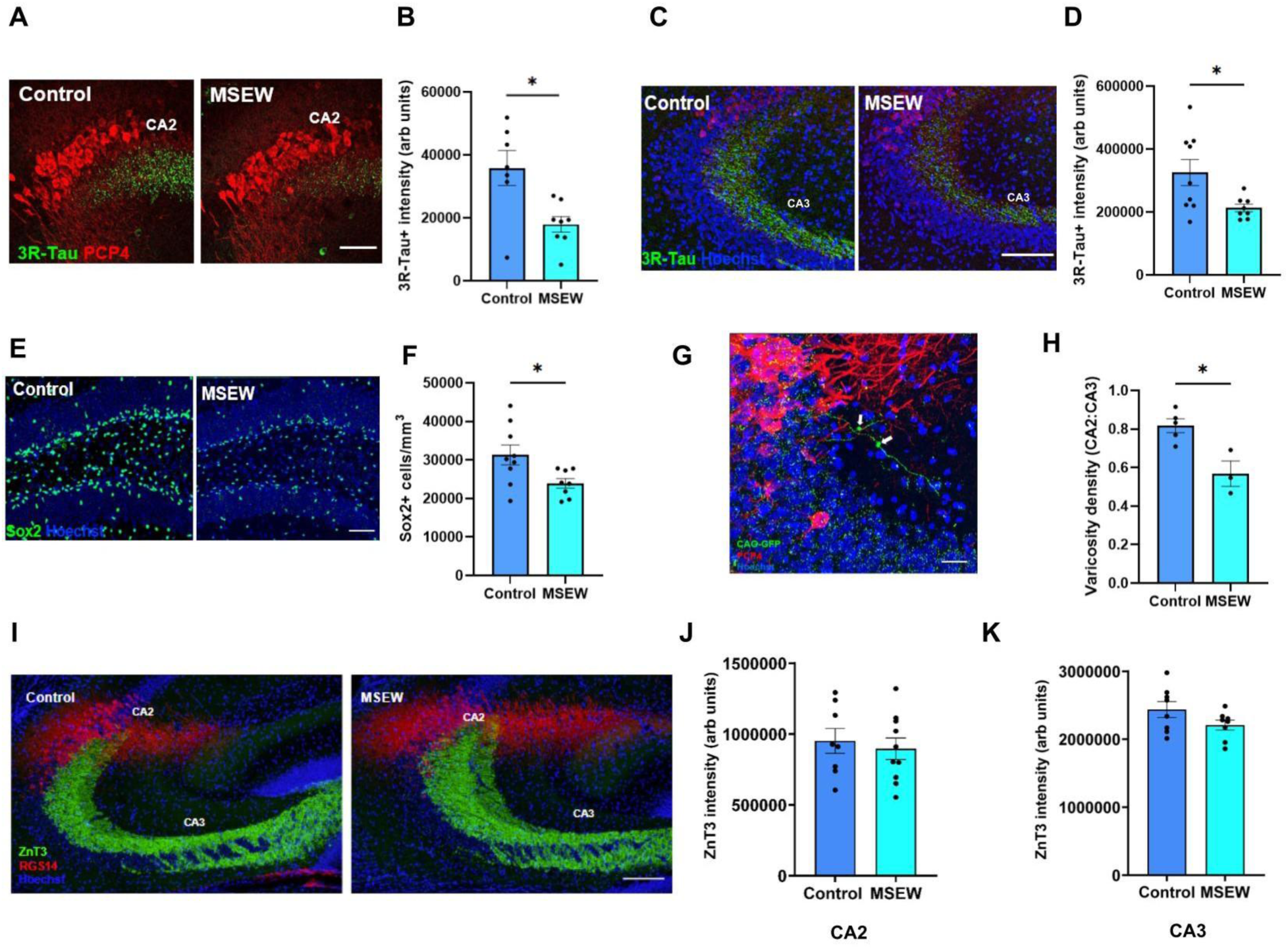
MSEW males have altered progenitor cells and adult born granule cell projections. **A)** Confocal images of 3R-Tau+ staining (green) in dorsal CA2 labeled by PCP4 (red) of control-reared and MSEW males. Scale bars = 40 μm. **B)** MSEW males showed a significant reduction in 3R-Tau+ staining in CA2 compared to control-reared animals. **C**) Confocal images of 3R-Tau+ staining (green) in dorsal CA3 of control-reared and MSEW males. Scale bars = 60 μm. **D**) MSEW males showed a significant reduction in 3R-Tau+ staining in CA3 compared to control-reared animals. **E**) Confocal images of Sox2+ cells (green) in dorsal dentate gyrus of control-reared and MSEW males. Scale bars = 40 μm. **F**) MSEW males had a significant reduction in Sox2+ cells in dorsal dentate gyrus compared to control-reared animals. **G**) Confocal image of GFP+ axons (green) and varicosities (white arrows) in the CA2 labeled by PCP4 (red). Scale bars = 20 μm. **H**) MSEW males had significantly lower proportion of CA2:CA3 GFP+ varicosities than control-reared males. **I**) Confocal images on ZnT3+ staining (green) in the CA3 and CA2 labeled with RGS14 (red) of Control-reared and MSEW males. Scale bars = 60 μm. **J)** Control-reared and MSEW males had similar intensities of ZnT3+ labeling in dorsal CA2. **K)** Control-reared and MSEW males had similar intensities of ZnT3+ labeling in dorsal CA3. *p<0.05. n=7-10 for immunolabeling studies; n=3-5 for retroviral labeling studies. Abbreviations: maternal separation with early weaning (MSEW), zinc transporter 3 (ZnT3).

**Figure S2:**
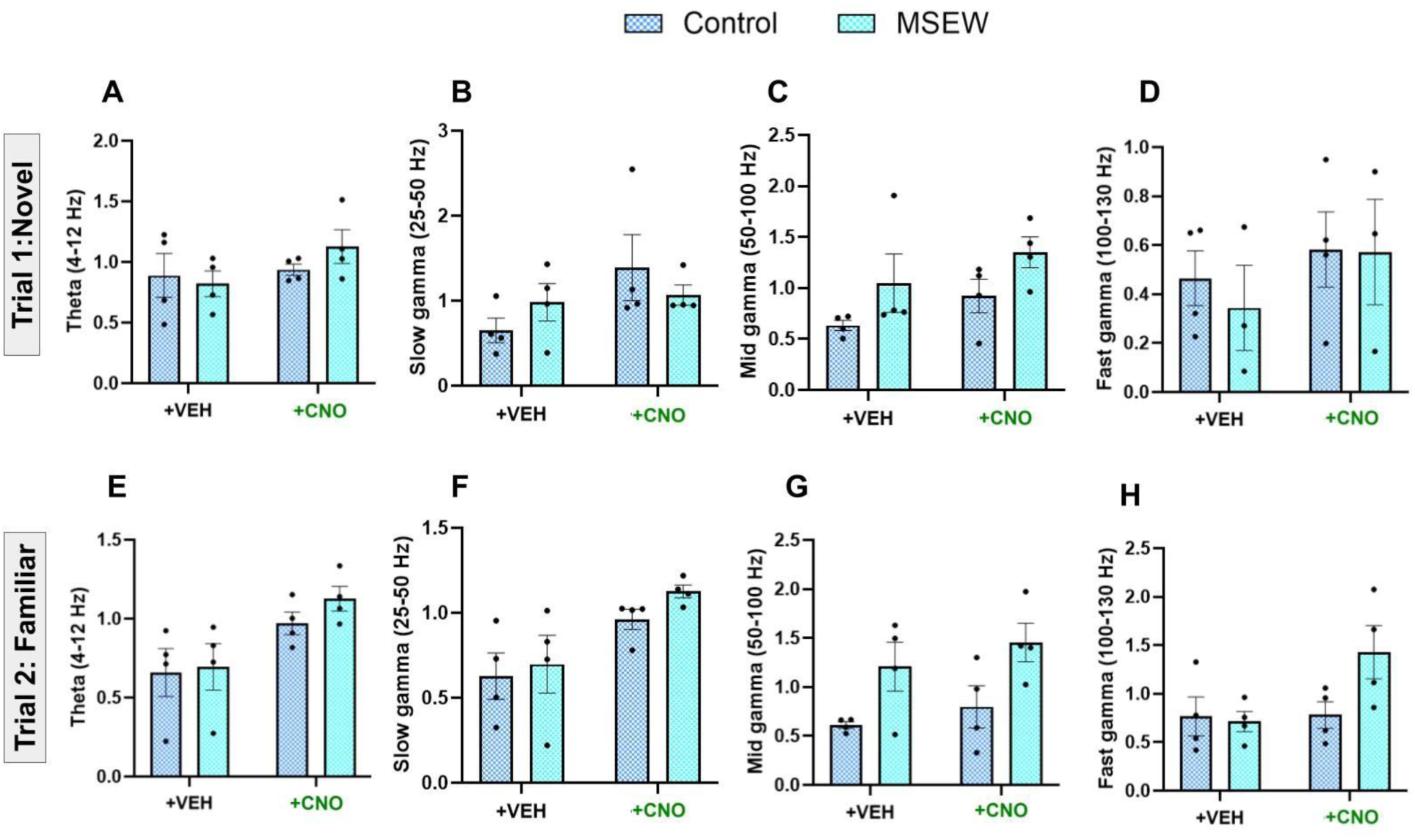
MSEW males show no detectable differences in CA2 theta or gamma power from control-reared males. No differences in CA2 theta (**A**), slow gamma (**B**), mid gamma (**C**) or fast gamma (**D**) power were observed in trial 1 between control-reared and MSEW males with vehicle or CNO treatment. No differences in CA2 theta (**E**), slow gamma (**F**), mid gamma (**G**) or fast gamma (**H**) power in Trial 2 were observed in trial 2 between control-reared and MSEW males with vehicle or CNO treatment. n=4 mice (4 electrodes per mouse). Abbreviations: maternal separation with early weaning (MSEW), vehicle (VEH)

**Figure S3:**
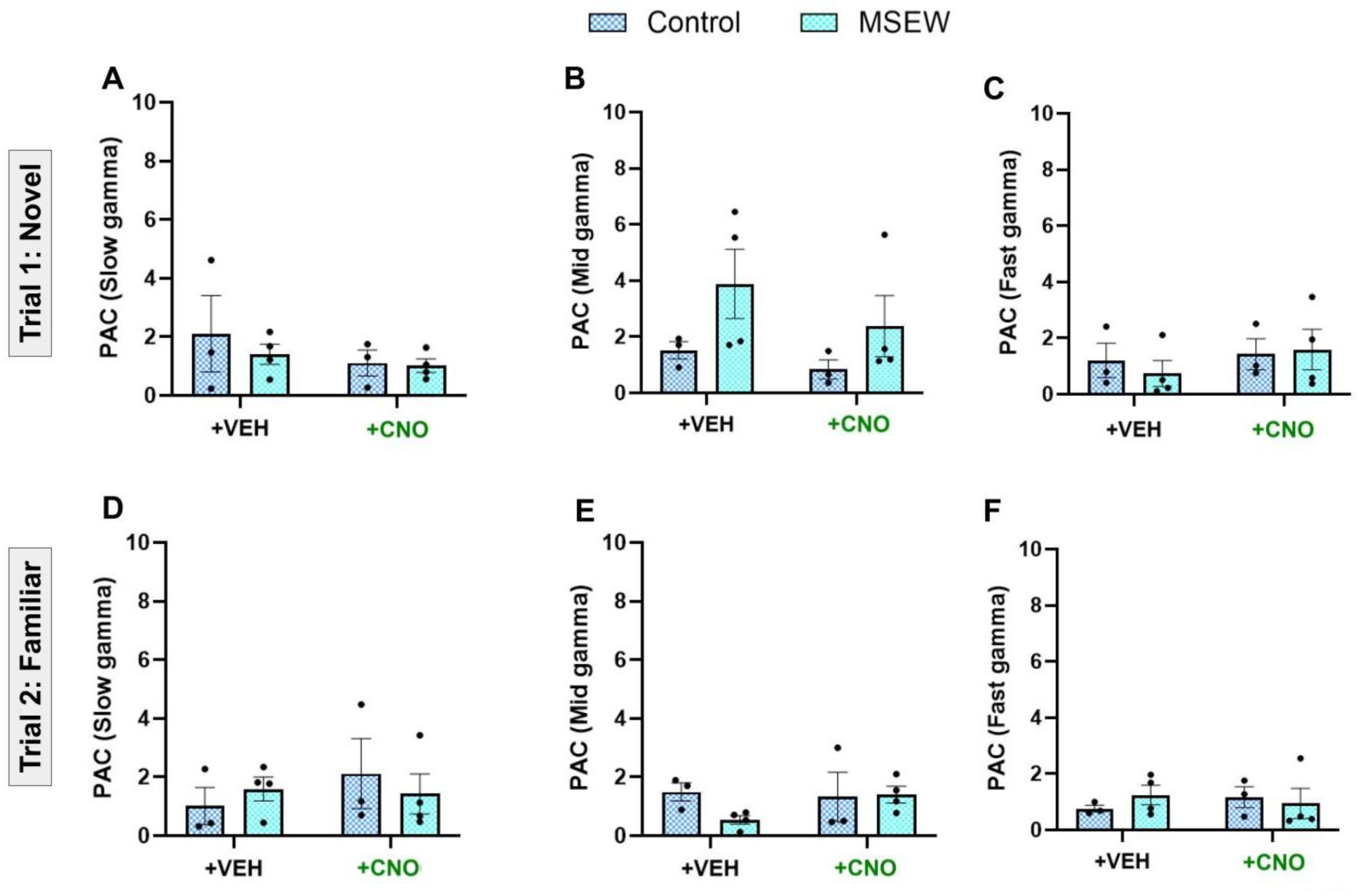
MSEW males show no detectable differences in CA2 phase amplitude coupling from control-reared males. No differences in CA2 theta-slow gamma (**A**), theta-mid gamma (**B**), or theta-fast gamma (**C**) PAC were observed in trial 1 between control-reared and MSEW males with vehicle or CNO treatment. No differences in CA2 theta-slow gamma (**D**), theta-mid gamma (**E**) or theta-fast gamma (**F**) PAC were observed in trial 2 between control-reared and MSEW males with vehicle or CNO treatment. n=3-4 mice (4 electrodes per mouse). Abbreviations: maternal separation with early weaning (MSEW), vehicle (VEH), phase-amplitude coupling (PAC).

**Figure S4:**
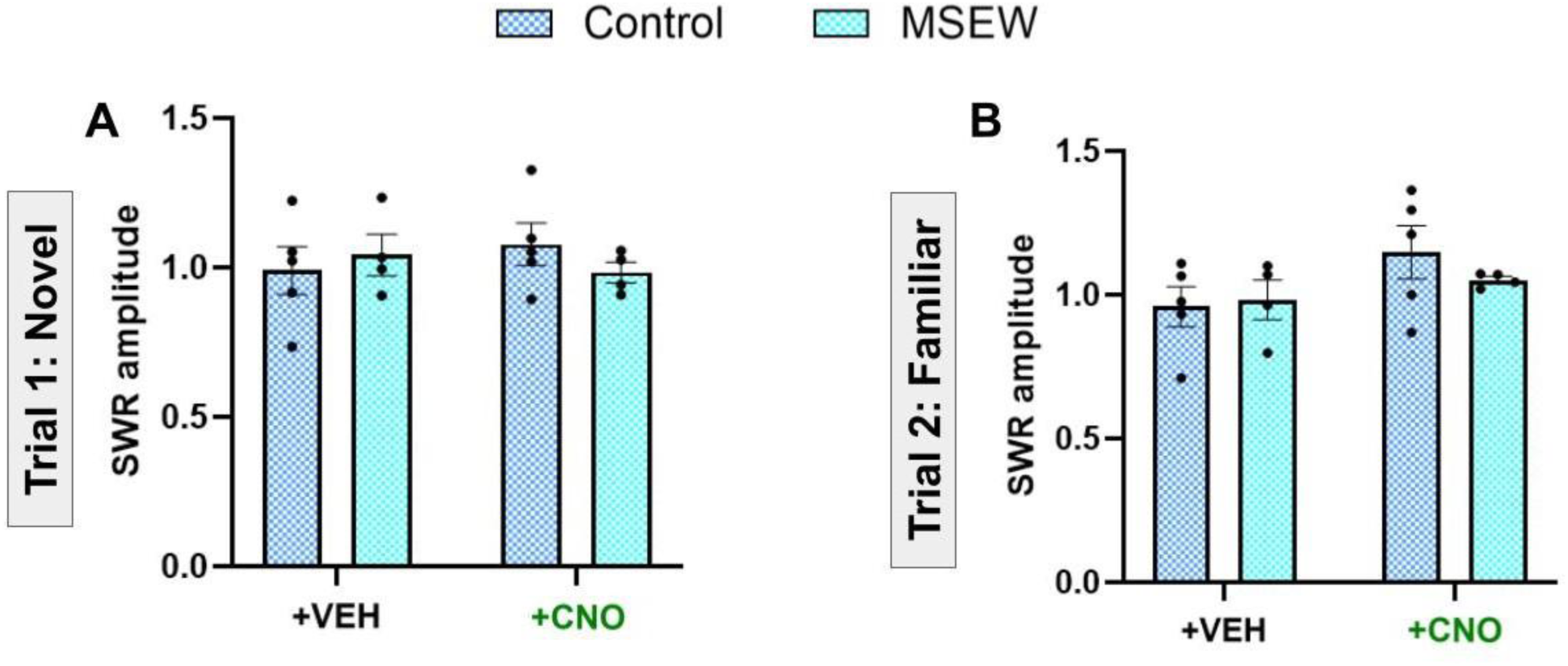
MSEW males show no detectable differences in SWR amplitude from control-reared males. No differences in CA2 SWR amplitude in trial 1 (**A**) or trial 2 (**B**) were observed between control and MSEW males with vehicle or CNO treatment. n=3-4 mice (4 electrodes per mouse). Abbreviations: maternal separation with early weaning (MSEW), sharp wave ripples (SWR), vehicle (VEH

## Notes

### Competing Interest Statement

The authors have declared no competing interest.

